# Manufacturing DNA in E. coli yields higher fidelity DNA than *in vitro* enzymatic synthesis

**DOI:** 10.1101/2023.09.12.557453

**Authors:** Steven J. Hersch, Siddarth Chandrasekaran, Jamie Lam, Nafiseh Nafissi, Roderick A. Slavcev

## Abstract

The rise of biotechnologies such as gene therapy have brought DNA vectors to the forefront of pharmaceutical development. The quality of the genetic starting material plays a pivotal role in determining the quality of the final product. In this study we examined the fidelity of DNA replication using enzymatic methods (*in vitro*) compared to plasmid DNA produced *in vivo* in *E. coli*. Next-generation sequencing approaches predominantly rely on *in vitro* polymerases, which have inherent limitations in sensitivity. To address this challenge, we introduce a novel assay based on loss-of-function (LOF) mutations in the conditionally toxic *sacB* gene. Our findings show that DNA production in *E. coli* results in significantly fewer LOF mutations (approximately 80-to 3000-fold less) compared to various enzymatic DNA synthesis methods. This includes the most accurate PCR polymerase (Q5) and a commonly employed rolling circle amplification (RCA) DNA polymerase (Phi29). These results suggest that using low-fidelity starting material DNA synthesized *in vitro* by PCR or RCA may introduce a substantial number of impurities, potentially affecting the quality and yield of final pharmaceutical products. In summary, our study underscores that DNA synthesized *in vitro* has a significantly higher mutation rate than DNA produced traditionally in *E. coli*. Therefore, utilizing *in vitro* enzymatically-produced DNA in biotechnology and biomanufacturing may entail considerable fidelity-related risks, while DNA starting material derived from *E. coli* substantially mitigates this risk, enhancing overall quality in the production processes.

## Introduction

Plasmids, along with other DNA vectors such as minicircles and linear derivatives, play a central role as starting materials, intermediates, drug substances, and drug products in the manufacturing of various therapeutics. These applications encompass DNA and mRNA vaccines, gene editing templates, and gene therapies employing both viral and non-viral vectors.Globally, over 400 million people suffer from rare diseases and could benefit from gene therapies^1^. However, high prices and complication risks are hurdles that must be overcome before gene therapy can reach its full potential^2,3^. Using high quality materials in manufacturing could increase potency (lower doses), improve production yield and efficiency (reduce prices), and reduce the potential risk of complications due to impurities in the product.

Recently, there has been increasing interest in the use of *in vitro* synthesized DNA. DNA made by polymerase chain reaction (PCR) or rolling circle amplification (RCA) can be produced rapidly and without the requirement of a living cell or cell bank. Nonetheless, even though the polymerase enzymes used in these systems are advertised as high fidelity, a lingering question persists: can they uphold the same degree of accuracy as DNA produced in a living organism?

To avoid damaging mutations during reproduction, all living organisms have evolved mechanisms to ensure that DNA is copied correctly. This replication fidelity begins with accurate base selection by DNA polymerase, followed by proofreading capability where high-fidelity polymerases reverse their progression to remove and replace incorrect bases^4^. DNA polymerases used *in vitro* for PCR or RCA-based DNA production include these two mechanisms^5^. However, in addition to base selection and proofreading, living cells such as *E. coli* that have been used for decades in the manufacturing of pharmaceutical products, have additional mechanisms to ensure replication fidelity. Most prominently is mismatch repair (MMR), which recognizes mismatched bases produced on the unmethylated daughter DNA strand, excises the inaccurate copy, and replaces it with the correct sequence dictated by the methylated parental strand^6,7^. This process improves the sequence accuracy of DNA production *in vivo* by approximately an additional 1000-fold^4,8–10^.

In this work we aimed to directly compare the sequence fidelity and quality of DNA starting material produced by *in vivo* (*E. coli*) and *in vitro* (DNA polymerase) procedures. Polymerases are traditionally tested by replicating *lacZ*, cloning it into a plasmid vector, and using blue/white screening to identify inactivating mutations in *lacZ*^11^. However, the sensitivity of this test is limited; at high fidelity the few white colonies cannot be accurately identified among the blue background. We also could not use next-generation sequencing (NGS) since NGS relies on *in vitro* polymerases with inherent error rates comparable to those used for *in vitro* DNA synthesis^5,12^. Therefore, the sensitivity of NGS is limited to about 0.1% mutations (assuming Q30 quality data) and any fidelity benefits of *in vivo* production would be masked by the error rate of the NGS enzyme^12^.

To overcome constraints posed by existing methodologies, we introduce an innovative and highly sensitive approach to assess the accuracy of DNA synthesis. Our ‘Sucrose Toxicity’ (SuTox) fidelity assay leverages positive selection for loss-of-function (LOF) mutants, effectively eliminating background and achieving exceptional sensitivity. Using the SuTox assay, we provide compelling evidence of the superior fidelity associated with DNA produced *in vivo* compared to DNA synthesized by *in vitro* enzymatic techniques.

## Results

### Design of the Sucrose Toxicity (SuTox) fidelity assay

We designed a screen for loss-of-function (LOF) mutations that generate a quantifiable phenotype in bacterial colonies. We chose the conditionally lethal gene, *sacB*, as our template for replication. SacB is harmless when *E. coli* are grown in standard media, avoiding selective pressure during *in vivo* replication. However, in the presence of sucrose, the SacB protein (levansucrase) produces a compound (levan) that accumulates in the periplasm and is toxic to *E. coli*^13^. Therefore, faithful replication of *sacB* prevents bacterial growth on agar plates containing sucrose. Colonies with LOF mutations in *sacB* survive, allowing simple visualization and quantification. This approach allows us to plate billions of bacteria without significant background, increasing sensitivity.

We employed a construct containing *sacB* and chloramphenicol resistance (chloramphenicol acetyl transferase) genes (Figure 1A). We cloned this cassette into a precursor plasmid used to produce double-stranded linear covalently closed DNA minivectors (msDNA™)^14,15^. Additionally, we added inverted terminal repeats (ITRs) to flank each end of the cassette to assess if these problematic palindromic sequences – used in rAAV manufacturing – would influence replication fidelity^16,17^. We replicated the ITR-*sacB-cat*-ITR cassette *in vivo* (by amplifying in *E. coli*) or *in vitro* using PCR or RCA DNA polymerization reactions (Figure 1B). We then cloned the synthesized DNA into a pUC19 vector, transformed highly competent cells, and plated on selective agar with or without sucrose. We selected for both the *sacB-cat* insert cassette (chloramphenicol) and the pUC19 vector (ampicillin). Notably, the ampicillin also counter-selects for the starting plasmid template, eliminating potential background from any carryover during cloning. Since survival on sucrose implies a LOF mutation(s) in the *sacB* gene or its expression, colonies growing in the presence of sucrose can be quantified to assess the accuracy of DNA production processes.

**Figure 1:**
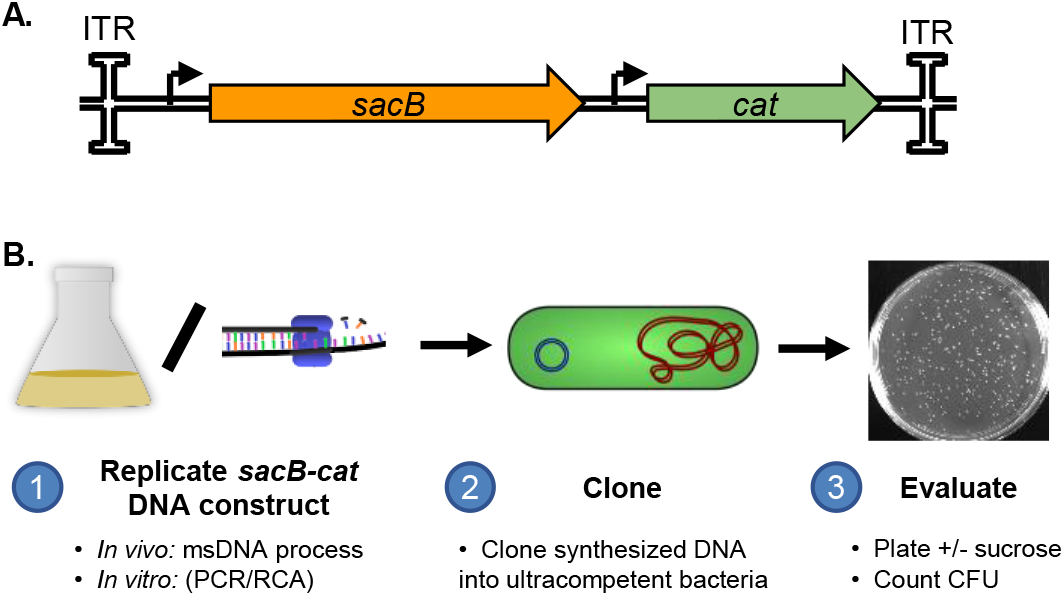
Measuring DNA replication fidelity by the Sucrose Toxicity (SuTox) assay. **A**. Overview of the template DNA construct. Chloramphenicol resistance (chloramphenicol acetyl transferase; *cat*) allows for selection of successful transformants. The SacB protein is toxic to bacteria in the presence of sucrose, allowing for positive selection of mutants: Faithful replication of the *sacB* gene results in bacterial cell death whereas if *sacB* is mutated (LOF) during DNA synthesis then a colony will grow on sucrose. Palindromic ITRs were included on the ends to investigate if they influence replication accuracy. **B**. Overview of the SuTox process. First, the *sacB-cat* DNA construct is replicated *in vivo* in bacterial cells or *in vitro* using PCR or rolling circle amplification (RCA). Second, the product is cloned into a vector and transformed into ultracompetent cells. Third, the transformation is plated with or without sucrose to quantify *sacB* mutants or total transformants (respectively).

### DNA manufactured in E. coli has higher fidelity than any enzymatic method tested

Using the SuTox fidelity assay, we compared the accuracy of DNA synthesis by multiple

*E. coli* strains (*in vivo*) and three different DNA polymerases (*in vitro*). We observed far fewer colonies on sucrose plates with *E. coli* transformed by DNA replicated *in vivo* in *E. coli* compared to any of the three enzymatic *in vitro* processes tested (Figure 2A). We quantified *sacB* mutants (CFU on sucrose), total transformants (CFU without sucrose), and the number of DNA doublings during synthesis (see Materials & Methods). By normalizing *sacB* mutants to total transformants and DNA doublings, we were able to accurately calculate mutation rates for each synthesis method. Compared to *in vivo* replication strain MBI3, we observed 2983, 737, and 84-fold higher mutation rates for Taq, Phi29, and Q5, respectively (Figure 2B). Notably, the same experiment conducted with an ITR-free *sacB-cat* cassette yielded similar data (Figure S1), suggesting that ITRs did not influence gene of interest (GOI) fidelity. We included a commonly used *E. coli* strain, BL21, as a benchmark for comparison to the MBI strains (Figure S1).

**Figure 2:**
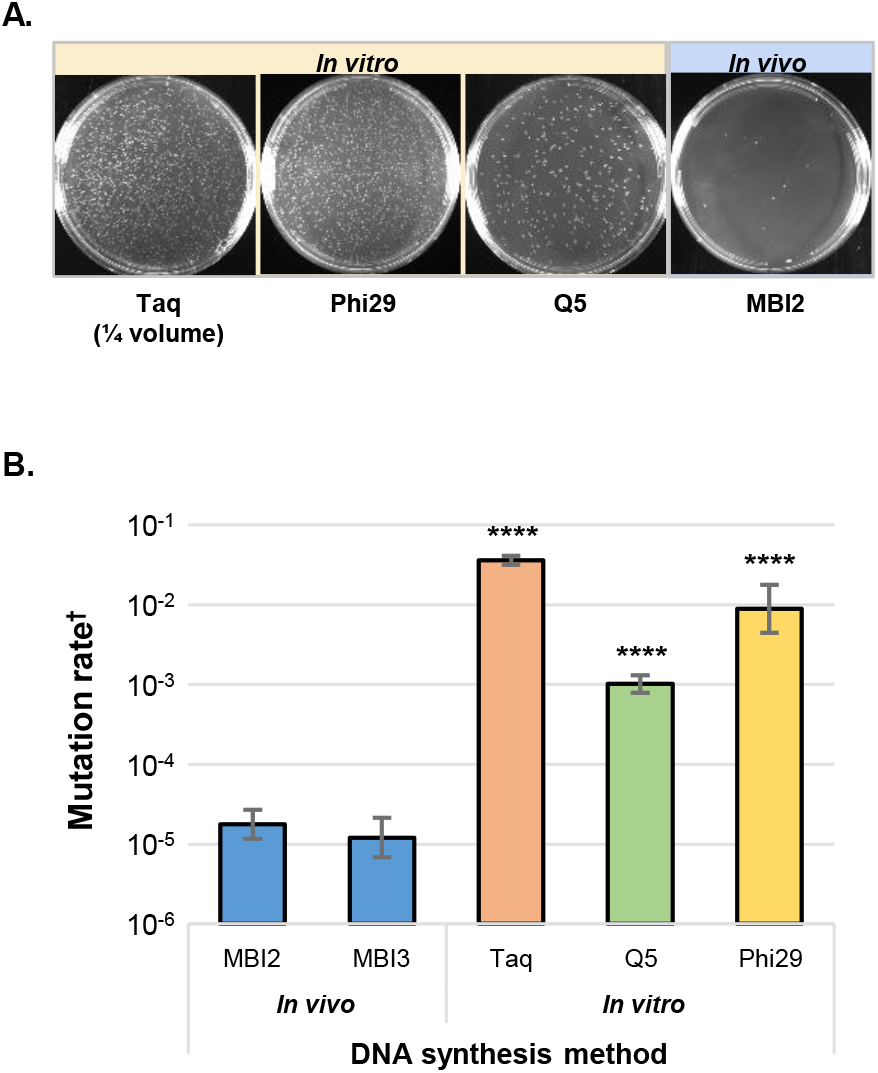
*DNA manufactured in* E. coli *has fewer LOF mutations than any enzymatic method tested*. **A**. Representative images of sucrose plates with transformations of *sacB-cat* DNA generated by PCR (Taq and Q5), RCA (Phi29), or *in vivo* in *E. coli*. For Taq, ¼ of the volume was plated relative to the other images. **B**. LOF mutation rates identified using the SuTox method with a *sacB-cat* cassette containing inverted terminal repeats (ITRs). DNA was synthesized *in vivo* (in two different *E. coli* strains) or *in vitro* (PCR with Taq or Q5 polymerases, or RCA with Phi29 polymerase). Bars show the average of three biological replicates and error bars show one standard deviation. One-way ANOVA with Dunnett’s test (compared to MBI2): ****, p < 0.0001. † As described in the text, mutation rate was calculated as *sacB* mutants / total transformants / DNA doubling.

### In vivo DNA production reduces mutation rate per kilobase by several orders of magnitude

We inferred that bacterial growth on media containing sucrose could be attributed to a LOF mutation occurring within the *sacB* open reading frame (ORF) or its associated promoter, encompassing a total length of 1533 bp. Due to the uncertainty surrounding the specific mutation sites responsible for LOF mutations, we calculated the mutation rate per nucleotide incorporated by dividing mutation rate by the entire 1533 bp. Our data revealed that replicating DNA in the MBI3 *E. coli* strain yields an estimated LOF error rate of approximately 7.9x10^-9^ / bp, or 0.00079% per kb (Table 1). For context, Phi29 (applied in RCA of DNA) showed an error rate of 0.58% per kb. Notably, even Q5, advertised as the highest fidelity PCR polymerase^5^, is better than Phi29 but still exhibited a significantly higher error rate compared to any of the *E. coli* strains tested.

**Table 1:** Approximate LOF mutation rates per kilobase synthesized.

| DNA synthesis method |  | Mutation rate / kb ‡ |
| --- | --- | --- |
| *In vivo* | MBI2 | 0.0012% |
|  | MBI3 | 0.00079% |
| *In vitro* | Taq | 2.3% |
|  | Q5 | 0.066% |
|  | Phi29 | 0.58% |

‡ As described in Materials and Methods, mutation rate / kb was calculated as: *SacB* mutation rate (from SuTox data in Figure 2), divided by the length of the *sacB* ORF and promoter (1533 bp), multiplied by 1000 bp / kb, multiplied by 100%.

## Discussion

In this work we have introduced the SuTox fidelity assay for the sensitive quantification of loss-of-function (LOF) mutations in DNA, whether generated *in vivo* or *in vitro*. The assay’s sensitivity primarily hinges on ligation efficiency and competency of the transformed cells. In one instance, a replicate using MBI2 fell below the detection limit (BDL), yielding no colonies on the sucrose plate. The other replicates for MBI2 were comparable to other *in vivo* strains, yielding few but quantifiable CFU on sucrose plates. The BDL replicate contributes to a lower average, but larger variance (Figure S1). The detection limit is also influenced by the number of DNA replication cycles. Since *in vivo* samples originated from a single transformed plasmid whereas the enzymatic methods started with nanograms of template DNA, the *in vivo*-produced DNA underwent additional replication cycles before assessment, enhancing their detection limit.

Importantly, the SuTox fidelity assay does not detect mutations that do not impede SacB activity (e.g. silent mutations). Additionally, as not all possible LOF sites within *sacB* are known, we calculated mutations per nucleotide using the entire length of the promoter and *sacB* ORF rather than solely the LOF subset. Consequently, the overall mutation rate is likely higher than the LOF mutation rate. This holds true across all tested conditions and the relative comparisons between DNA synthesis methods remain accurate. Given that the impact on biotechnology primarily stems from mutations affecting functionality, the LOF mutation rate is a more relevant metric of assessment.

The data obtained using the SuTox fidelity assay closely aligns with mutation rates documented in existing literature: Particularly, our mutation rate per kb per doubling data (Table 1) are similar to findings using SMRT sequencing for Q5^5^ and radioactive nucleotide misincorporation for Phi29^18^. Interestingly, our assessment reveals that Taq polymerase shows five-fold higher fidelity compared to previously reported data^5,11^. This discrepancy may stem from potential improvements in enzyme or buffer conditions offered by different Taq polymerase vendors. Furthermore, prior studies have explored *in vivo* DNA replication through long-term passaging of *E. coli* and monitoring changes in chromosomal DNA^9,10,19–21^. These works have demonstrated even higher fidelity for *in vivo* DNA replication than what we present here. While this could partly be attributed to variations in mutation rates between plasmid and genomic DNA, it is more likely attributable to distinctions in detection limits between the SuTox assay and long-term evolution assays, which involve more DNA replication cycles and employ a large screening template (chromosomal DNA). However, it is important to note that the SuTox fidelity assay offers the advantage of directly comparing replication fidelity between *in vivo* and *in vitro* methods, using plasmid templates commonly used in biotechnology. In this direct comparison, the difference in fidelity of *in vivo* over *in vitro* replication (approximately 2-3 orders of magnitude) aligns well with previous findings that quantified the benefit of MMR^4,8–10^, lending further credence to our results.

To illustrate the practical significance of these findings, consider an rAAV Cis (ITR-GOI-ITR) construct that is 4.7 kb long. Companies using Phi29 to manufacture their rAAV construct could expect that approximately 2.7% of their final GOI products contain LOF mutations and more may have mutations with potential consequences other than LOF of the GOI. These nonfunctional but packaged impurities could potentially increase immunogenicity and reduce dose potency of the final rAAV product. Synthesis of rAAV also requires two additional constructs, ‘RepCap’ (5.5 kb) and ‘Helper’ (13 kb). Using DNA synthesized with Phi29, the odds of a triply transfected cell having no LOF mutations in any of the three constructs is only 86.5%, compared to 99.98% using DNA produced in *E. coli*. Similarly, consider a large nonviral gene therapy construct expressing the 12kb dystrophin gene. Companies using Phi29 could expect approximately 7% of their products to be defective due to a LOF mutation, compared to only about 0.01% for companies using DNA produced *in vivo* in *E. coli*.

Finally, besides the occurrence of LOF mutations, there exists the potential for dominant negative gain-of-function mutations in certain instances. To illustrate, consider a gene therapy scenario involving expression of the tumour suppressor gene p53. P53 has numerous known dominant negative mutations that can promote cancerous growth^22,23^. Drawing from our LOF data, it becomes evident that the likelihood of encountering dominant negative mutations and their associated complications is remarkably higher when using DNA starting material synthesized using enzymatic *in vitro* methods.

Current guidelines for manufacturing plasmids and other DNA materials primarily address their use as therapeutic agents^24^. However, these DNA forms (including traditional plasmids, miniplasmids, and linear DNA vectors) also serve as pivotal starting materials in the production of viral vectors and mRNA. As regulatory requirements become increasingly stringent, it becomes imperative to consider diverse quality and testing needs to harmonize DNA production and procurement practices. Depending on the intended role of DNA in drug development, whether as a starting material, intermediate, drug substance, or drug product, various regulatory agencies could introduce distinct quality and fidelity standards.

In summary, using the SuTox assay to directly compare replication fidelity, we have established that DNA produced *in vivo* in *E. coli* cells significantly surpassed the accuracy of PCR or RCA methods. While *in vitro* methods remain invaluable for generating new clonal constructs, industrial scaling up to quantities necessary for drug production may introduce cumulative mutation risk. Employing *E. coli*-based *in vivo* generated DNA, such as plasmid or ministring DNA (msDNA) can effectively reduce mutations, thereby mitigating risk and enhancing the overall quality of the final product.

## Materials & Methods

### Bacterial strains and plasmids

All bacterial strains were *E. coli* derivatives. MBI strains could not be disclosed but a commonly used lab strain, BL21, was also tested for comparison. The starting plasmid, containing the ITR-*sacB-cat*-ITR construct (and the version without ITRs) were initially generated in Mediphage’s proprietary precursor plasmid for manufacturing ministring DNA (msDNA), a linear double-stranded DNA molecule with covalently closed ends^14,15^. msDNA is bacterial-sequence-free but is produced by an *in vivo* process originating from plasmid DNA. Following replication, the cassette was inserted into pUC19 for transformation on sucrose plates.

### DNA synthesis by in vivo or in vitro methods

For *in vivo* DNA replication, we transformed the starting plasmid into *E. coli* strains and grew the transformations overnight on selective LB agar plates. Single colonies were inoculated into 2 mL liquid LB cultures, grown shaking for 16h at 37 °C, then treated with a commercial miniprep kit (OmegaBioTek D6945-02) to obtain *in vivo-*synthesized plasmid DNA. Since each colony originates as a single bacterium transformed with a single molecule of plasmid, replication from the starter plasmid began from that single plasmid molecule. Therefore, the DNA input was calculated as the mass of one plasmid molecule. The DNA output was calculated as the concentration of the miniprep (obtained using a nanodrop spectrophotometer) multiplied by the elution volume.

For *in vitro* replication, we designed primers binding just outside of the *sacB-cat* construct that added either a SacI or SalI restriction enzyme site. Using these primers, we amplified the cassette by PCR using Taq (Froggabio T-500) or Q5 (NEB M0491) polymerases, applying their respective manufacturer-suggested buffers and thermocycling conditions. The same primers were used to guide RCA with Phi29 polymerase (NEB M0269). Following PCR or RCA, the enzymes and buffer reagents were removed using a commercial PCR purification kit (Thermo Fisher K0702) to obtain *in vitro*-synthesized DNA. Since Phi29 produces multimers that are difficult to purify, we digested the completed Phi29 reaction with SacI and SalI restriction enzymes prior to the PCR purification protocol. The DNA input was calculated as the quantity of starter plasmid added to the reaction as template multiplied by the length of the amplified region as a fraction of the total plasmid size. The DNA output was calculated as the concentration of the PCR purification (obtained using a nanodrop spectrophotometer) multiplied by the elution volume.

### Transformation of synthesized DNA and calculation of mutation rate

The *in vivo* and *in vitro* synthesized DNA were digested with SacI and SalI restriction enzymes (NEB R3156, R3138), as was a pUC19 vector with ampicillin resistance. We used a commercial gel extraction kit (Thermo Fisher K0691) to isolate the *sacB-cat* fragment and ligated each insert:vector combination overnight with T4 ligase (NEB M0202). We also included a no-insert reaction as a negative control. The ligations were transformed into high efficiency (1-3 × 10^9^ CFU/μg pUC19 DNA) competent cells (NEB C3040). The transformations were serially diluted and plated on LB with ampicillin (100 μg/ml), chloramphenicol (25 μg/ml), and either 1% NaCl (standard Miller LB) or 6% sucrose. The plates were grown at 37 °C for 16-24 h.

Colony forming units (CFU) were counted for each sample with and without sucrose. Any CFU counted from the negative control transformation were subtracted as background for all samples. Samples with no CFU on sucrose after background subtraction were treated as below detection limit (BDL) and calculated as if 0.5 CFU were on the sucrose plate. CFU from the sucrose plates were counted as *sacB* mutants. CFU on standard LB plates were counted as total transformants. The number of DNA doublings was calculated as Log_2_(DNA output / input). Finally, the mutation rate was calculated as the *sacB* mutants / total transformants / DNA doubling. Paraphrased, this calculates the fraction of DNA molecules that contain a LOF *sacB* mutation, normalized to how many times the original template was replicated.

The mutation rate calculated from the SuTox method describes LOF mutations in the *sacB* gene. To obtain an estimate of the number of mutations per bp replicated, we divided the mutation rate by the length of the *sacB* ORF and promoter (1533 bp). We then multiplied by 1000 bp to get mutations / kb replicated, followed by multiplying by 100% to express the values as a percentage.

## Data Availability Statement

Data will be made available on request.

## Acknowledgements

The authors thank Dr. Ko Currie for helpful discussions, and Dr. Tao Dong and Dr. William Navarre for their donation of plasmids. Funding for this work was supported by Mediphage Bioceuticals, Inc. and the National Sciences and Engineering Council of Canada (grant #391457).

## Author Contributions

SJH designed the SuTox fidelity assay, performed and analyzed the experiments, and prepared the manuscript and figures. SC contributed to the SuTox assay design. SC and JL assisted with experiments. NN and RAS supervised the study. All authors contributed to manuscript revision.

## Supplementary Figure Legends

**Figure S1:**
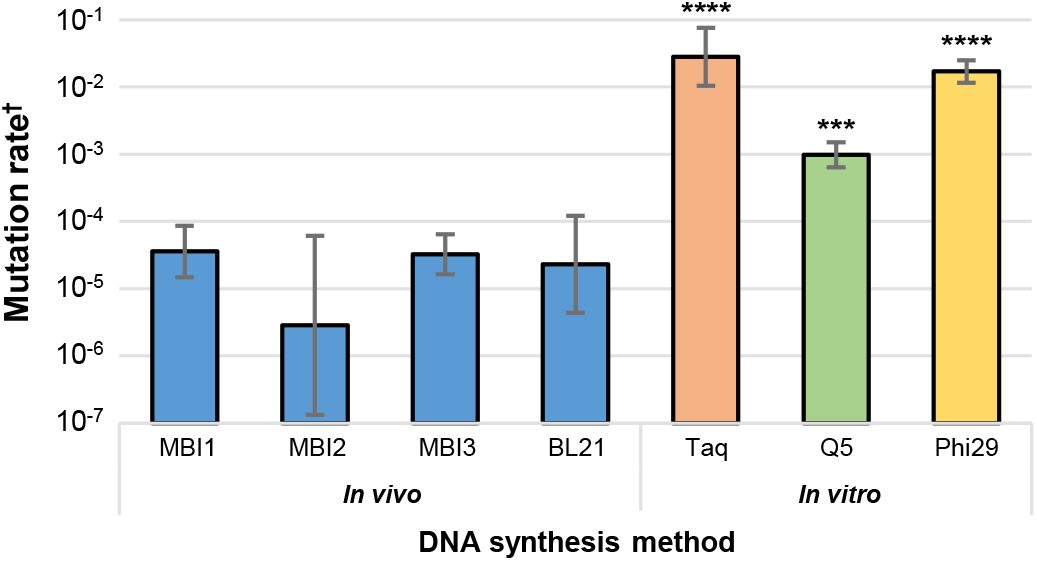
LOF mutation rates identified using the SuTox method with a *SacB-cat* cassette that does *not* contain ITRs. DNA was synthesized *in vivo* (in four different *E. coli* strains) or *in vitro* (PCR with Taq or Q5 polymerases, or RCA with Phi29 polymerase). Bars show the average of at least three biological replicates and error bars show one standard deviation. One-way ANOVA with Dunnett’s test (compared to MBI2): ***, p < 0.001; ****, p < 0.0001. † As described in the text, mutation rate was calculated as *sacB* mutants / total transformants / SDNA doubling.

